# *Prochlorococcus* phage ferredoxin: structural characterization and electron transfer to cyanobacterial sulfite reductases

**DOI:** 10.1101/2020.02.07.937771

**Authors:** Ian J. Campbell, Jose L. Olmos, Weijun Xu, Dimithree Kahanda, Joshua T. Atkinson, Othneil N. Sparks, Mitchell D. Miller, George N. Phillips, George N. Bennett, Jonathan J. Silberg

**Affiliations:** Biochemistry and Cell Biology Graduate Program, Rice University, 6100 Main Street, MS-140, Houston, TX 77005, USA; Department of Biosciences, Rice University, 6100 Main Street, MS-140, Houston, TX 77005, USA; Department of Chemistry, Rice University, 6100 Main Street, MS-140, Houston, TX 77005, USA; Department of Chemical and Biomolecular Engineering, Rice University, 6100 Main Street, MS-362, Houston, TX 77005, USA; Department of Bioengineering, Rice University, 6100 Main Street, MS-142, Houston, TX 77005, USA

**Keywords:** cyanobacteria, electron transfer (ET), ferredoxin (Fd), marine, bacteriophage, sulfite reductase (SIR), reductase, hydrogen sulfide

## Abstract

Marine cyanobacteria are infected by phage whose genomes encode ferredoxin (Fd) electron carriers. While these Fds are thought to redirect the energy harvested from light to phage-encoded oxidoreductases that enhance viral fitness, it is not clear how the biophysical properties and partner specificities of phage Fds relate to those in photosynthetic organisms. Bioinformatic analysis using a sequence similarity network revealed that phage Fds are most closely related to cyanobacterial Fds that transfer electrons from photosystems to oxidoreductases involved in nutrient assimilation. Structural analysis of myovirus P-SSM2 Fd (pssm2-Fd), which infects *Prochlorococcus marinus*, revealed high similarity to cyanobacterial Fds (≤0.5 Å RMSD). Additionally, pssm2-Fd exhibits a low midpoint reduction potential (−336 mV vs. SHE) similar to other photosynthetic Fds, albeit lower thermostability (T_m_ = 28°C) than many Fds. When expressed in an *Escherichia coli* strain with a sulfite assimilation defect, pssm2-Fd complemented growth when coexpressed with a *Prochlorococcus marinus* sulfite reductase, revealing that pssm2-Fd can transfer electrons to a host protein involved in nutrient assimilation. The high structural similarity with cyanobacterial Fds and reactivity with a host sulfite reductase suggest that phage Fds evolved to transfer electrons to cyanobacterial-encoded oxidoreductases.

## Introduction

*Prochlorococcus marinus* is thought to be the most prevalent photosynthetic organism on Earth, with a global abundance of 10^27^ cells (1), making it a key player in biogeochemical processes. This oligotrophic cyanobacterium inhabits the euphotic zone of oceans (40°N to 40°S) and is projected to expand both in density and range as global temperatures rise (1). Collectively fixing four gigatons of carbon annually (1), the diverse ecotypes of *P. marinus* thrive at many depths, where light, nutrient availability, and temperature vary (2–4). Up to 24% of the CO_2_ fixed by *P. marinus* is released as dissolved organic carbon, providing critical feedstocks for heterotrophic organisms and supporting the larger oceanic ecosystem (5).

The core metabolic systems in *P. marinus* have diverged significantly from other cyanobacteria due in part to genomic streamlining (6–10). Additionally, *P. marinus* lacks phycobilisomes widely used by other cyanobacteria for light harvesting and instead uses a divinyl chlorophyll a_2_/b_2_ complex (11). Furthermore, this cyanobacterium contains only one set of core photosystem components while other cyanobacteria maintain several homologs (8, 12). Nitrogen assimilation pathways have been pruned, with species often lacking the ability to reduce either nitrate or nitrite (13, 14). Various ecotypes are incapable of heterotrophic growth, lacking several tricarboxylic acid cycle genes (8).

The life cycle of *P. marinus* is influenced by viruses. Cyanophages that infect *P. marinus* have evolved genomes with as many as 327 open reading frames (ORFs) (15). Viruses from three clades infect *P. marinus*, including T4-like myoviruses, T7-like podoviruses, and, less commonly, members of *Siphoviridae* (16). In some ecosystems, as many as 50% of cyanobacteria may be infected at any point in time (17). Some cyanophages exhibit infection cycles that are closely tied to day-night rhythms of cyanobacteria, although individual viruses vary in their responses (18). Once infection begins, many host genes are repressed in favor of phage homologs (19), which can decrease cyanobacterial carbon fixation, potentially keeping ∼5 gigatons of carbon in the atmosphere (20). While recent estimates suggest that cyanophages are limited by energy rather than nutrients (21), our understanding of their controls over host electron transfer (ET) remains limited.

Following infection, cyanophages express metabolic genes to modulate host photosynthesis (22). Phage can express their own photosystem II reaction core (D1 protein) during infection (17, 23, 24), with viral transcripts equaling or even exceeding host transcripts in some settings. Certain cyanophages encode an entire photosystem I (PSI) complex that is thought to be a more promiscuous oxidizer of electron carriers than host PSI, which may push host cells toward cyclic photosynthesis (25). Viral plastocyanin homologs, which possess altered charged surfaces compared with host plastocyanin, can also be expressed and are thought to transfer electrons directly to cytochrome aa_3_ oxidase rather than PSI as a mechanism for managing redox balance (26). Other phage genes implicated in oxygenic photosynthesis include *hli* (high-light inducible protein), *pebA* (15,16- dihydrobiliverdin:Fd reductase), *psbD* (PSII D2 protein), *pcyA* (phycocyanobilin:Fd reductase), and *petF* (Fd) (15). Although the sequence similarities of these genes to their cyanobacterial homologs give clues to their cellular roles, we do not know how these proteins interact with host proteins to tune the host electron fluxome.

The *Prochlorococcus* P-SSM2 phage Fd (pssm2- Fd) supports the reduction of phage phycoerythrobilin synthase (PebS), which catalyzes the reduction of biliverdin IX_α_ to *3E/3Z* phycoerythrobilin (27, 28). While prior studies showed that some Fds support ET to diverse partner oxidoreductases (29–32), it remains unclear how the partner specificity of this and other phage Fds compare to host Fds, which can function as ET hubs that couple light harvesting to a range of reductive metabolic reactions (31–33). In addition, it is not known how the primary structure and biophysical properties of phage Fds compare to host Fds.

In this study, we used a sequence similarity network (SSN) analysis to examine how cyanophage and cyanobacterial Fds relate to one another. We also determined the first structure of a cyanophage Fd, which has high structural similarity to cyanobacterial Fds. We show that this Fd has a low midpoint potential characteristic of photosynthetic Fds, establish that this protein has a low thermostability (T_m_ = 28°C), and show that it supports ET to a host oxidoreductase involved in sulfur assimilation. These results suggest that viral Fds support ET to cyanobacterial-encoded oxidoreductases, and they extend our understanding about the ways viruses interact with cyanobacterial hosts.

## Results

### Cyanophage and host Fd bioinformatics

To better understand the prevalence of cyanophage Fds, we searched all 8392 viral genomes available in the Integrated Microbial Genomes and Microbiomes database for ORFs matching the Interpro signature for [2Fe-2S] Fds (34). This genome mining yielded 26 Fds from phages that have been shown to infect *Prochlorococcus* and *Synechococcus* (including pssm2-Fd), a Fd from an uncultured marine virus, and a Fd from the giant *Pseudomonas* phage OBP (35). All viral ORFs encoded Fds with 95 to 97 residues, with the exception of the *Pseudomonas* phage OBP which was 83 residues.

Sequence similarity networks (SSNs) represent a simple way to visualize how protein paralogs relate to one another in their primary structure, and they provide a frame of reference for establishing how phage-encoded proteins may have evolved from host proteins. Additionally, the individual clusters in these networks often include protein homologs with similar functional properties (36), so they can provide hints about the possible functions of phage-encoded proteins. To understand which cyanobacterial Fds are most closely related to phage Fds, we used an SSN to compare viral Fds with 807 cyanobacterial Fds obtained from a recent genome mining effort (37). *Zea mays* Fd (zm-Fd1) was included as a frame of reference because this protein has been intensively studied (38, 39). When the Fds were compiled into a SSN (40), the cyanobacterial Fds presented over a dozen distinct clusters, which arose because of variation in Fd length (Figure 1A). Across the twelve largest clusters, the average Fd length varies by ∼100 amino acids. These clusters also vary in the number of cyanobacterial species represented and the average Fd size, with the top five most species diverse clusters being VI (100 ±8 residues) > IX (115 ±9 residues) > VII (106 ±13 residues) > I (79 ±2 residues) > XI (163 ±8 residues) (Dataset S1).

**Figure 1.**
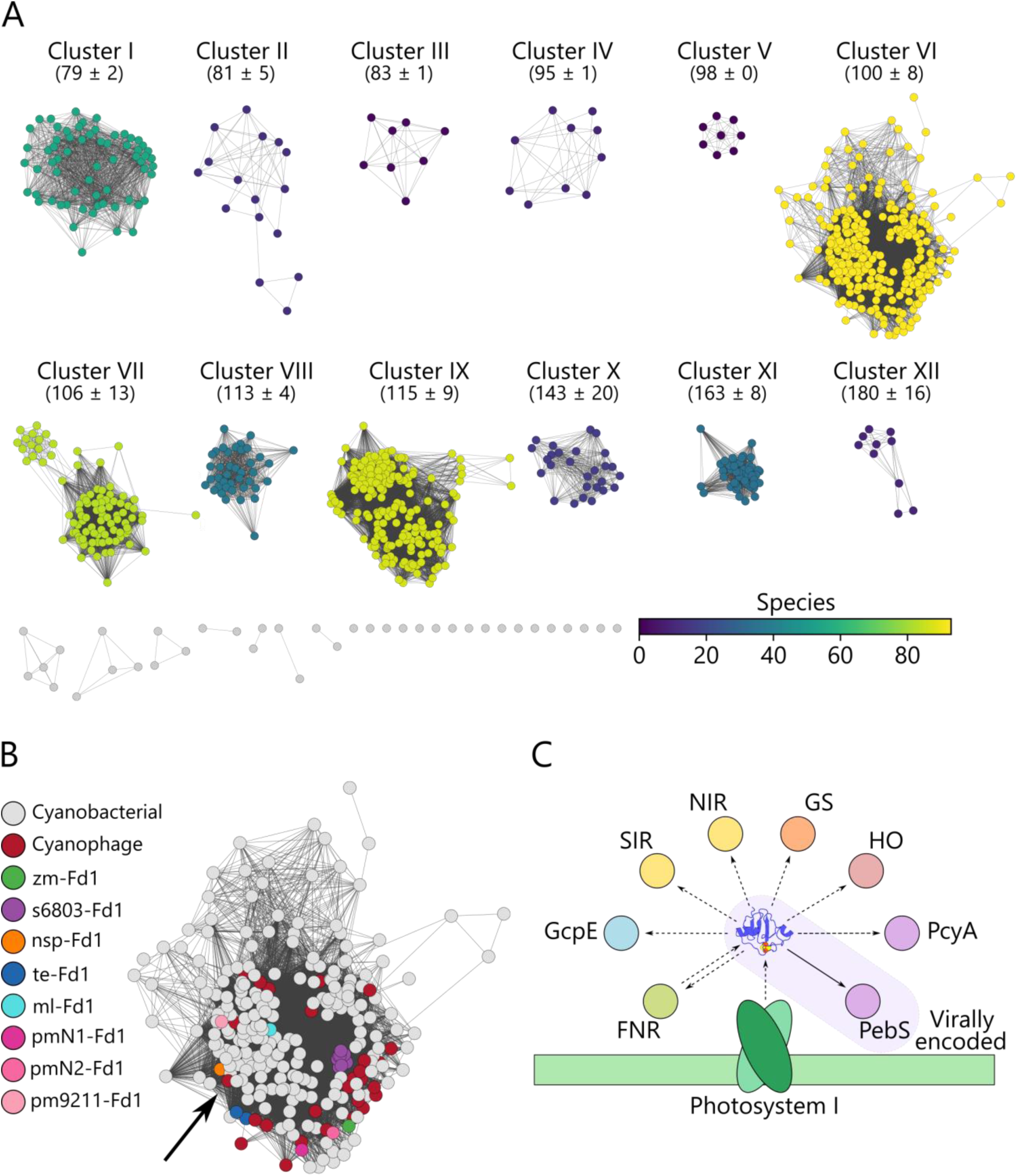
Network analysis of cyanophage and cyanobacterial Fds. (**A**) A network illustrating the relationships of 807 cyanobacterial and 28 cyanophage Fds. Edges are drawn between Fd pairs that exhibit a sequence alignment score ≥30. For the largest clusters (I to XII), the average protein lengths encoded by the ORFs in the cluster are provided ±1σ. Clusters are colored using a viridis gradient by the number of species represented in each cluster. (**B**) Cyanophage and cyanobacterial Fds are highlighted within Cluster VI with zm-Fd1 as a frame of reference. The arrow indicates pssm2-Fd. Fds from multiple species and strains share the same sequence as s6803-Fd1 and te-Fd1, so they are all colored identically. (**C**) *P. marinus* partner oxidoreductases that have the potential to interact with pssm2-Fd during an infection. Abbreviations: Fd (ferredoxin), Fd-NADP reductase (FNR), 4-Hydroxy-3-methylbut-2-enyl diphosphate synthase (GcpE), sulfite reductase (SIR), nitrite reductase (NIR), glutamate synthase (GS), heme oxygenase (HO), phycocyanobilin:Fd oxidoreductase (PcyA), and PebS (phycoerythrobilin synthase).

The phage Fds all localize to Cluster VI in close association with cyanobacterial Fds (Figure 1B), with the exception of *Pseudomonas* phage OBP Fd, which does not cluster with any Fds. Cluster VI includes numerous structurally-characterized Fds, including Fds from *Synechocystis sp*. PCC 6803 (s6803-Fd1), *Nostoc sp*. PCC 7119 (nsp-Fd1), *Thermosynechococcus elongatus* (te-Fd1), and *Mastigocladus laminosus* (ml-Fd1). Additionally, this cluster contains zm-Fd1, the Fd included as a frame of reference.

Among the Fds in Cluster VI, interactions with a range of oxidoreductases have been documented. s6803-Fd1 binds photosystem I (PSI), glutamate synthase (GS), and Fd-NADP reductase (FNR) (41–43). nsp-Fd1 interacts with FNR (44). te-Fd1 interacts with FNR, PSI, heme oxygenase (HO), PcyA, nitrite reductase (NIR), and 4-Hydroxy-3- methylbut-2-enyl diphosphate synthase (GcpE) (45–48). This latter oxidoreductase is in the non-mevalonate pathway for isoprenoid biosynthesis (47). The tight association of cyanophage and cyanobacterial Fds having a wide range of protein-protein interactions suggested cyanophage Fds might be able to support ET to host oxidoreductases. Given that sulfite reductase (SIR) and NIR have similar sequences, possess related folds, and utilize the same cofactors, we reasoned that cyanophage Fds could interact with SIR (38, 49, 50). pssm2-Fd is encoded by a myophage that infects *P. marinus*. By analogy to characterized host Fds, it may have the potential to interact with a total of eight host oxidoreductases (Figure 1C), in addition to the virally-encoded PebS (15, 27, 51).

### pssm2-Fd contains sequence features of host Fds

To better understand how pssm2-Fd structure relates to cyanobacterial Fds, we analyzed the pairwise sequence identity of pssm2-Fd and cyanobacterial Fds in *P. marinus* hosts that are infected by cyanophage P-SSM2 (51). Fds from three host strains were collected by protein BLAST and designated pmN1-Fd1 (ecotype NATL1A), pmN2-Fd1 (NATL2A), and pm9211-Fd1 (MIT9211) (52). Like pssm2-Fd, these Fds reside within Cluster VI. We also evaluated the pairwise identity of pssm2-Fd with structurally-characterized Fds, including cyanobacterial (s6803- Fd1, nsp-Fd1, te-Fd1, and ml-Fd1) and plant (zm- Fd1) Fds. This analysis revealed most (5/7) cyanobacterial Fds exhibit ≥70% identity with each other (Figure 2A), while pssm2-Fd exhibits a range of sequence identities with these Fds (40-60%). Interestingly, pssm2-Fd presents slightly higher sequence identities with Fds from non-host cyanobacteria than from host cyanobacteria. The Fds from *P. marinus* ecotypes NATL1A and NATL2A (pmN1-Fd1 and pmN2-Fd1) exhibit the lowest identity with pssm2-Fd (43%) and appear to be the two most divergent Fds in the set. pm9211- Fd1 exhibits higher sequence identity with pssm2- Fd (52%) and appears more closely related to other cyanobacterial Fds (s6803-Fd1, te-Fd1, nspFd1, and ml-Fd1) than the other *P. marinus* ecotypes.

**Figure 2.**
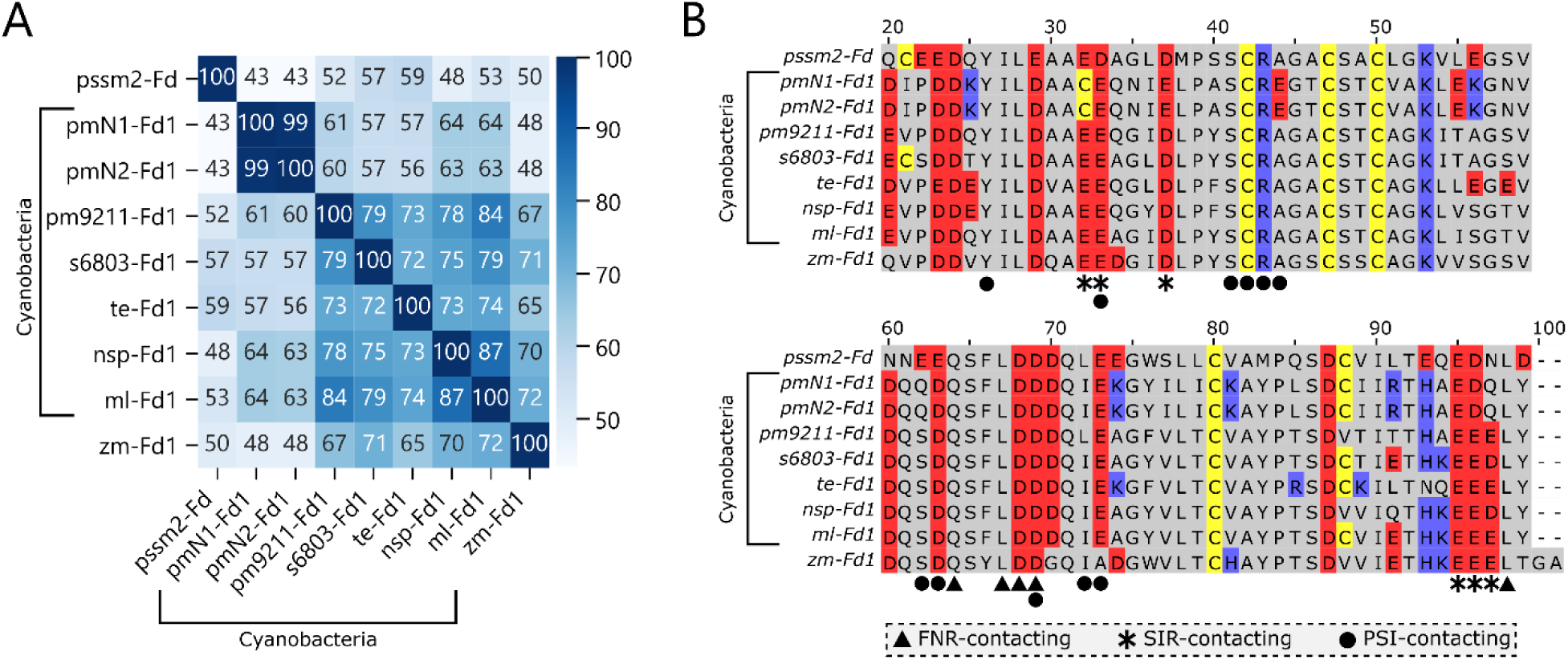
Cyanophage and host cyanobacterial Fd sequence comparisons. (**A**) A matrix showing the pairwise sequence identity of pssm2-Fd and Fds from host cyanobacteria, cyanobacteria with known structures, and one plant family member (zm-Fd1). The blue shading indicates the percentage sequence identity. (**B**) A multiple sequence alignment illustrates the conservation of cysteines (yellow), positively charged residues (blue), negatively charged residues (red), and residues implicated in binding FNR, SIR, and PSI (38, 39, 45). Residues are indexed with respect to ml-Fd1.

Fd binding specificities are influenced by charged surface residues and residues that make direct contacts with partner proteins (38, 39). To establish whether pssm2-Fd sequence is conserved at positions implicated in forming residue-residue contacts with partner proteins, a multiple sequence alignment was generated (Figure 2B). Residues implicated in contacting Fd partners from crystal structures of Fd-partner complexes are highlighted (38, 39, 45). In the case of the Fd-FNR complex (PDB = 1GAQ), all of the Fd residues that make contacts with FNR are completely conserved (39), suggesting a ubiquitous binding strategy to FNR across cyanobacteria. For the Fd-PSI complex (PDB = 5ZF0) and Fd-SIR (PDB = 5H92), greater variability is observed in the Fd residues that mediate partner contacts (38, 45). Only eight of the eleven Fd residues that contact PSI are either absolutely conserved or have residues with similar physicochemical properties. Similarly, four of the six Fd residues that contact SIR are either absolutely conserved or similar.

### pssm2-Fd is marginally stable

The tight association of pssm2-Fd with cyanobacterial Fds in Cluster VI suggested that these proteins may exhibit similar biophysical properties. To obtain recombinant pssm2-Fd for *in vitro* characterization, the gene encoding pssm2-Fd was overexpressed in *Escherichia coli* at 37°C and purified using a combination of anion exchange and size exclusion chromatography. This protocol yielded 0.6 mg/L of phage Fd, which was estimated to be >95% homogeneous by SDS-PAGE analysis. This yield is ∼25-fold lower than that obtained with a thermophilic cyanobacterial Fd using an identical expression protocol (53).

To determine if pssm2-Fd contains a [2Fe-2S] cluster, we acquired pssm2-Fd absorbance and circular dichroism spectra (Figures 3A, 3B). The absorption spectrum contains peaks (465, 420, and 330 nm) that are characteristic of holoFds (54, 55). Additionally, the circular dichroism spectrum presented ellipticity maxima (427 and 360 nm) and minima (550 and 505 nm) that are observed with holoFds (54, 55). The ratio of the ellipticity at 427 nm to pssm2-Fd concentration was comparable to a previously purified *Azotobacter vinelandii* Fd that had near stoichiometric cluster occupancy (56). This suggests that a significant fraction of pssm2- Fd contains a [2Fe-2S] cluster.

**Figure 3.**
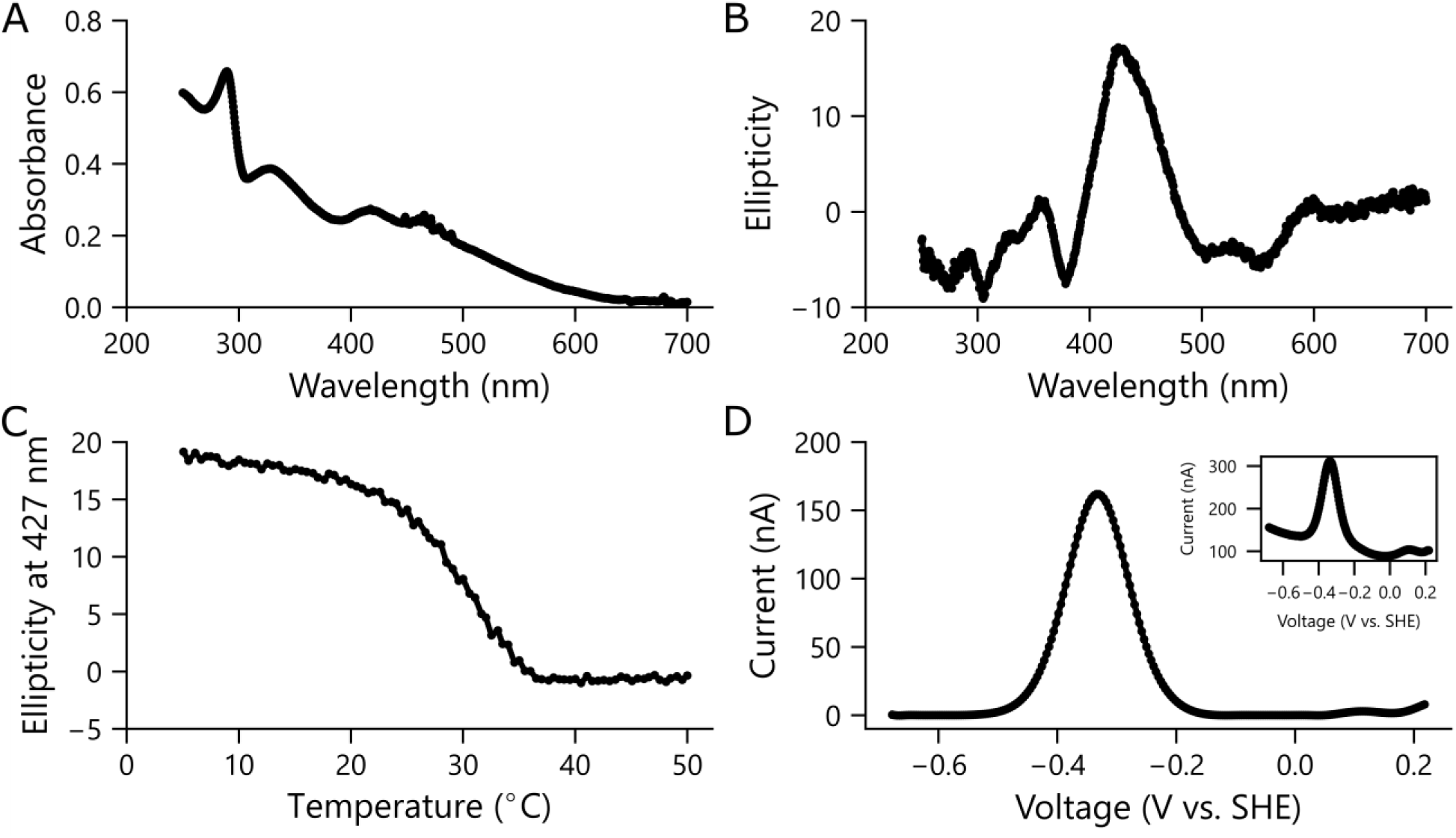
Biophysical properties of purified pssm2-Fd. (**A**) The absorbance and (**B**) circular dichroism spectra of purified pssm2-Fd (50 µM) dissolved in TED (25 mM Tris pH 8, 1 mM EDTA, and 1 mM DTT) reveals features characteristic of [2Fe-2S] Fds. (**C**). Effect of temperature on the ellipticity of pssm2-Fd (50 µM) in TED at 427 nm. The midpoint of the transition occurs at 29°C. (**D**) Square wave voltammetry of pssm2-Fd (620 µM) in 5 mM acetate, 5 mM MES, 5 mM MOPS, 5 mM TAPS, 5 mM CHES, 5 mM CAPS, and 100 mM NaCl (pH 7) reveals a midpoint potential of -334 mV. Voltammetry prior to background subtraction is shown as an inset.

Because the yield of pssm2-Fd was lower than a thermophilic Fd that we recently purified (53), we postulated that pssm2-Fd might have a lower melting temperature (T_m_). Iron-sulfur clusters often stabilize Fd structures, and in some cases only degrade after some secondary structure is lost (57, 58). As a [2Fe-2S] moiety is required for Fd electron transfer and cluster loss often leads to irreversible folding, we evaluated the effect of temperature on pssm2-Fd by measuring the ellipticity (at 427 nm) that arises from iron-sulfur cluster binding (Figure 3C) (59, 60). Assuming two-state unfolding (61), the fraction of folded pssm2-Fd at each temperature was analyzed in a Van’t Hoff plot. A linear fit of this plot yielded ΔH_unfolding_ and ΔS_unfolding_ (Figure S1), which were used to calculate T_m_ of 28°C.

To determine how the midpoint potential of pssm2- Fd relates to photosynthetic Fds, we characterized the electrochemical properties of this Fd using thin film, square wave voltammetry (62). With this analysis, pssm2-Fd presented a low midpoint potential (E_m_ = -336 mV) relative to a standard hydrogen electrode (Figure 3D), a value that is similar to other photosynthetic Fds.

### Cyanophage Fd structural characterization

We determined the pssm2-Fd crystal structure at 1.6 Å resolution. The resulting model (*R*_cryst_/*R*_free_ are 0.180/0.213) contains all 96 residues for each of the two monomers in the asymmetric unit, excluding the initiator methionines. The two subunits in the asymmetric unit superposition well, with a low root-mean-square deviation of all atomic coordinates (0.17 Å). The main source of variation between the chains arises from subtle differences within the beta sheets. While Chain A in the asymmetric unit is well ordered, Chain B has weaker density for residues around the iron-sulfur cluster with a poor fit to the electron density resulting in a large real-space R-value in this region. Therefore, non-crystallographic symmetry (NCS) restraints refinements were carried out using Chain A as a reference for Chain B. Additionally,

Chain A has density that is consistent with a [2Fe- 2S] cluster while the same region of Chain B has density that reflects a mixture of zinc and a [2Fe- 2S] cluster. This observation is likely due to the zinc present in crystallization conditions. Evidence in the literature indicates that zinc can interfere with and disrupt iron-sulfur clusters (63–66). To account for the poor density and the anomalous data, Chain B was modeled with grouped occupancies, where the model is a mixture of an intact cysteine-coordinated iron-sulfur cluster and a zinc ion with a hydration shell. An interpretation of these data is that as the cluster is disrupted, the surrounding residues become disordered. The crystallographic data collection and refinement statistics are provided in Table S1.

The structure of the pssm2-Fd is shown in Figure 4A. With five β-sheet strands and three α-helices, pssm2-Fd possesses a canonical β-grasp topology, which consists of a β-sheet that is four to five strands long with α-helices interspersed and packed against the sheet (67). This topology is observed in other Fds, such as ml-Fd1 and s6803-Fd1 (41, 68). To quantify how similar the pssm2-Fd is to cyanobacterial Fds in Cluster VI (1OFF, 5AUI, 1CZP, 1RFK, 3B2F), we calculated the pairwise RMSD between these structures and Chain A of pssm2-Fd (41, 68–71). The pssm2-Fd structure presents low RMSD (≤0.5 Å) in all atom positions with the cyanobacterial Fds (Table S2), including two thermophilic Fds (45, 68) and a plant Fd (zm- Fd1). Given the high similarity of Chain A to other Fd structures, we used this polypeptide for all subsequent structural analysis.

**Figure 4.**
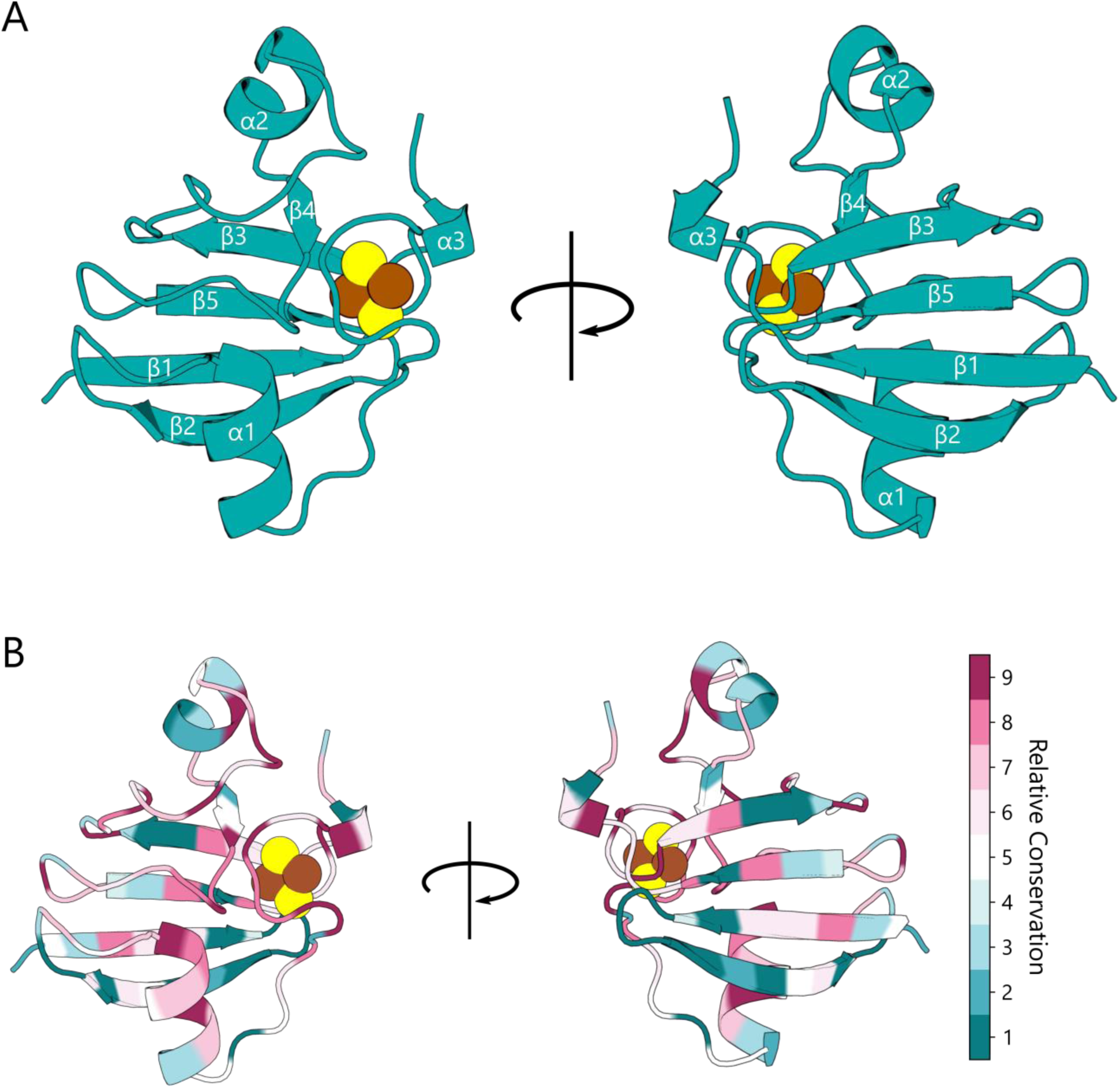
The phage Fd structure. (**A**) Front- and back-facing ribbon structures of the pssm2-Fd structure. Iron and sulfur atoms are shown as brown and yellow spheres, respectively. The α helices and β strands are labelled sequentially. (**B**) Ribbon structures of pssm2-Fd colored with ConSurf conservation scores generated using the Fds in Cluster VI. The most conserved residues are shown in magenta, while the least conserved residues are in cyan.

We next mapped the sequence variation of Fds within Cluster VI onto the pssm2-Fd structure (Figure 4B). To quantify variation, we calculated the conservation scores at each pssm2-Fd residue using ConSurf (72). This analysis revealed that the largest density of absolutely conserved residues occurs in the loop surrounding the [2Fe-2S] cluster, with the next most conserved regions being those adjacent to this loop. There are additional residues that are highly conserved across all of the Fds analyzed. However, these are dispersed across the structure and primarily localized to regions that are packed within the center of the pssm2-Fd structure or forming salt-bridges on the surface.

Analysis of the intramolecular interactions within pssm2-Fd and those observed within cyanobacterial Fds found in Cluster VI revealed additional differences (Table S2). Among the proteins analyzed, pssm2-Fd presents a lower number of salt bridges (n = 3) compared with all other Fds analyzed (n = 4 to 6). However, pssm2-Fd contains a similar number of total hydrogen bonds (n = 133) as the other Fds (n = 105 to 154). Only one salt bridge was conserved in all of the Fds, which arises from an interaction between an arginine at alignment position 43 and either a glutamate or aspartate situated between positions 29 and 32. This interaction may be conserved because it helps shape the [2Fe-2S] cluster-binding loop, as the arginine is directly adjacent to a cluster-ligating cysteine (Figure 2B).

### pssm2-Fd supports ET to a host SIR

In a previous study, we examined whether pssm2- Fd could support ET between *Zea mays* FNR (zm- FNR) and SIR (zm-SIR) using a synthetic ET pathway in *E. coli* that is required for cellular growth (53). In this assay, Fd-mediated ET from FNR to SIR is required for sulfur assimilation and growth on medium containing sulfate as the only sulfur source (Figure 5A) (73). This prior study found that pssm2-Fd could not support ET from zm-FNR to zm-SIR (53). However, these experiments were performed at 37°C where only a small fraction of pssm2-Fd is folded.

**Figure 5.**
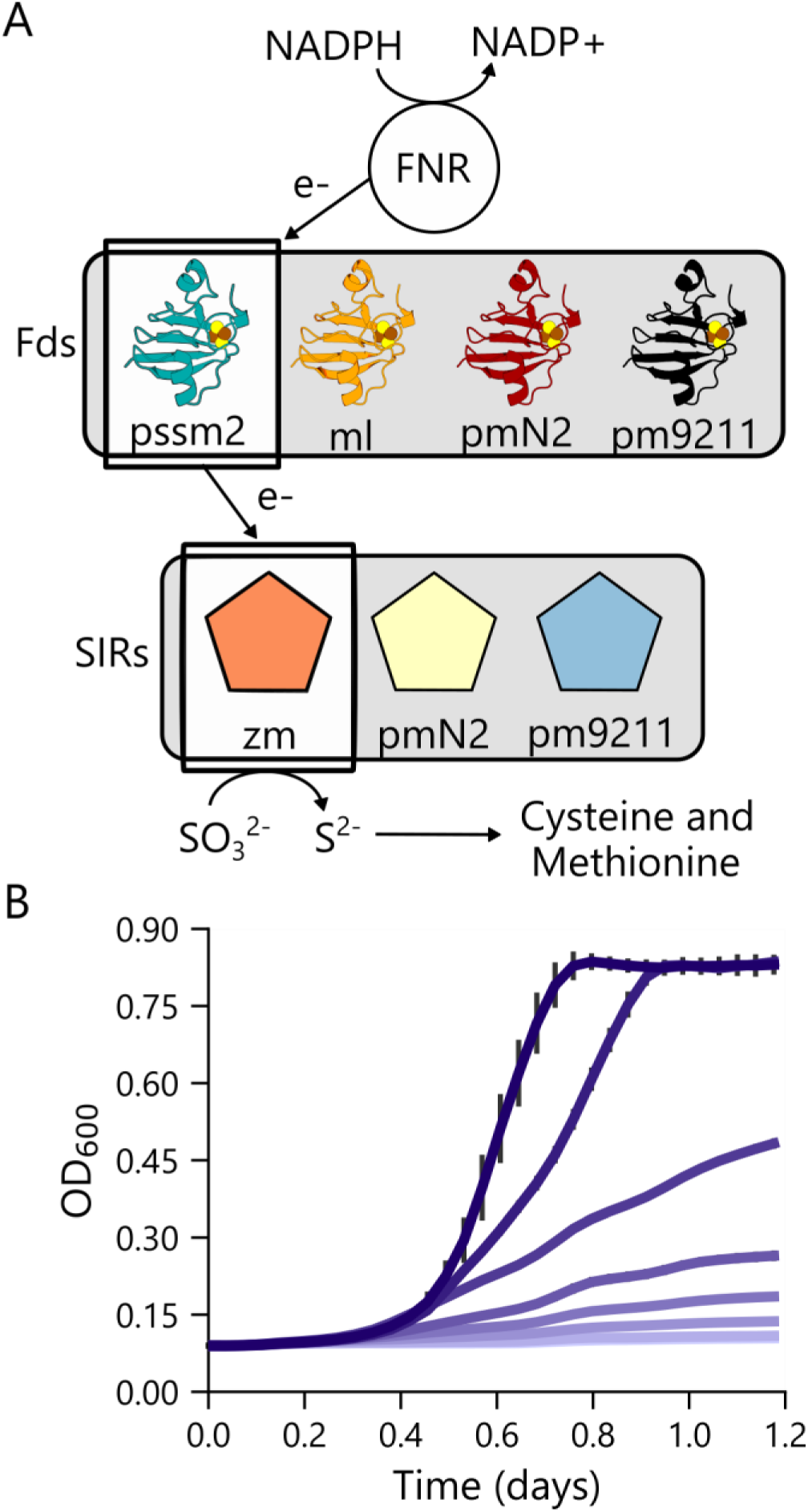
Monitoring pssm2-Fd ET using a cellular assay. (**A**) An *E. coli* auxotroph unable to produce sulfide can grow on sulfite as a sulfur source when cells express a three-component pathway consisting of FNR, Fd, and SIR. In this study, four different Fds and three different SIRs were examined. (**B**) The effect of aTc concentration on the growth complementation of *E. coli* EW11 transformed with vectors that express pssm2-Fd using an aTc-inducible promoter and zm-FNR/zm-SIR using constitutive promoters. The aTc concentrations (0, 1.6, 3.1, 6.3, 12.5, 25, 50, 100 ng/mL) are color coded as a gradient from light purple (no aTc) to dark purple (highest aTc). Average values from three biological replicates are shown with error bars representing ±1σ.

To increase the fraction of holo-pssm2-Fd in the cellular assay, we built an anhydrotetracycline (aTc)-regulated expression vector that initiates translation of pssm2-Fd using a synthetic ribosome binding site (RBS) with strong translation initiation. When the assay was performed using this vector, aTc-dependent complementation was observed at 37°C (Figure 5B). This finding indicates that pssm2-Fd supports ET from zm-FNR to zm-SIR but requires high protein expression to compensate for the low stability of pssm2-Fd.

To test whether pssm2-Fd can also support ET to host SIRs, we created vectors that constitutively express *P. marinus* NATL2A SIR (pmN2-SIR) and MIT9211 SIR (pm9211-SIR) in parallel with zm- FNR. SIR from the NATL1A host was omitted because it shares high sequence identity (98%) with pmN2-SIR (Figure S2). Analysis of the zm- Fd1/zm-SIR complex structure revealed that these *P. marinus* SIRs are identical with zm-SIR at five of the seven residues that bind zm-Fd1 (Figure S3) (38). We also built aTc-inducible vectors to express host Fds (pmN1-Fd1, pmN2-Fd1, and pm9211- Fd1). Using these vectors, we tested the ability of cyanophage and host Fds to support ET from zm- FNR to zm-SIR and from zm-FNR to two host SIRs (pmN2-SIR and zm-9211-SIR). Because ml-Fd1 supports ET from zm-FNR to zm-SIR (53), it was included in the analysis as a positive control. Assays were performed at 23, 30, and 37°C ±aTc, which regulates Fd expression.

At the highest temperature assayed (37°C), only cells expressing zm-SIR complemented growth when coexpressed with a Fd and FNR (Figure 6A). This growth was observed with ml-Fd1, pssm2-Fd, and pm9211-Fd1. Cells expressing pmN2-Fd did not support ET from zm-FNR to zm-SIR. At 30°C, complementation was again observed with the same three-component ET pathways (Figure 6B). Additionally, cells expressing pm9211-SIR complemented growth when coexpressing the zm- FNR and either the cognate host Fd, pssm2-Fd, or ml-Fd1. At 23°C, similar trends were observed with zm-SIR and pm9211-SIR as at 30°C (Figure 6C). Additionally, significant complementation, albeit small, was observed when pmN2-SIR was coexpressed with the pssm2-Fd and ml-Fd1.

**Figure 6.**
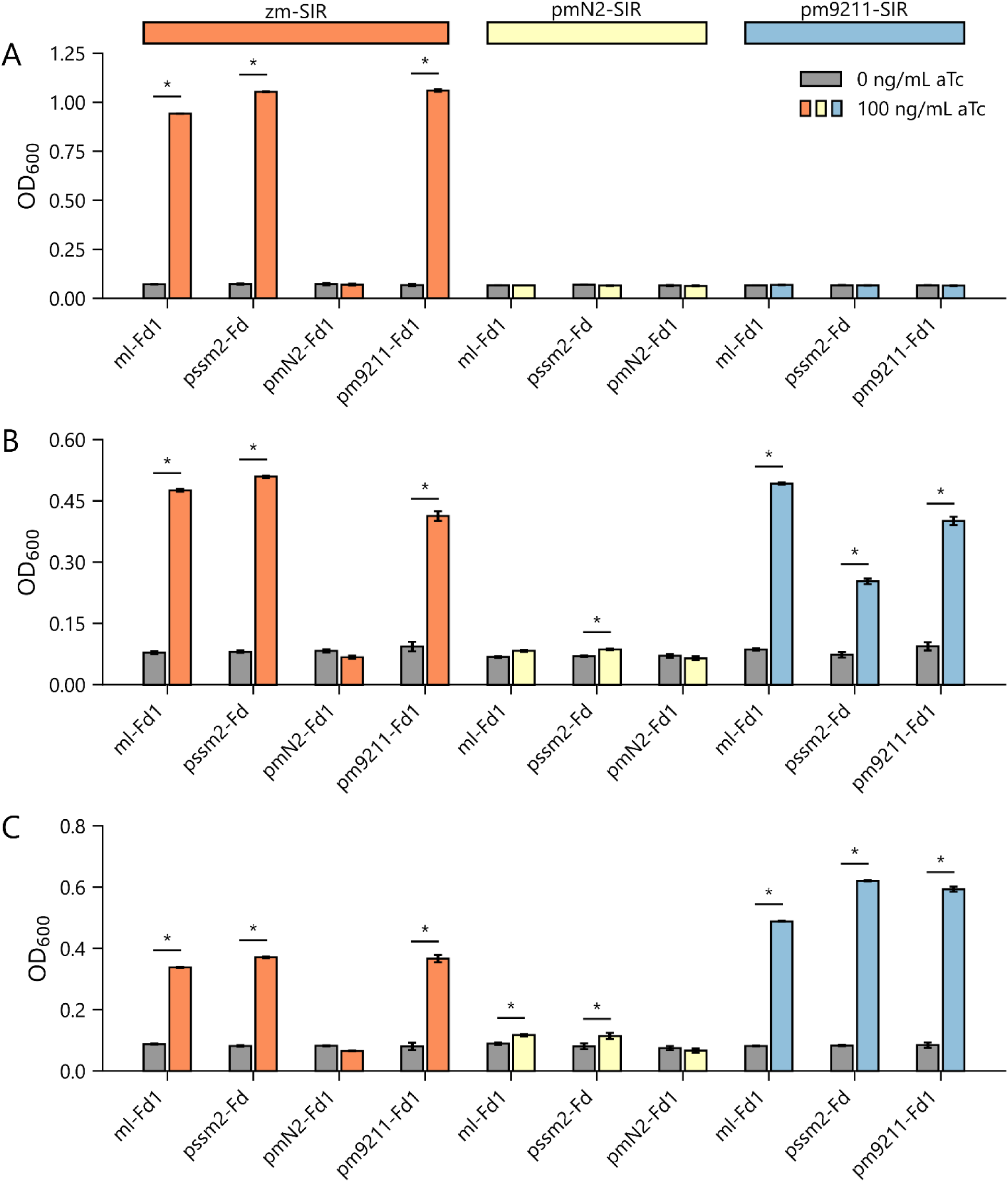
pssm2-Fd supports ET to host SIRs. Growth complementation observed at (**A**) 37°C, (**B**) 30°C, and (**C**) 23°C with *E. coli* EW11 transformed with vectors that constitutively express zm-FNR with three different SIRs (zm-SIR, pmN2-SIR, pm9211-SIR) and a vector that uses an aTc-inducible promoter to express different Fds (ml-Fd1, pssm2-Fd, pmN2-Fd1, pm9211-Fd1). Experiments were performed in the absence (grey bars) and presence (colored bars) of 100 ng/mL aTc to evaluate the dependence of growth upon Fd expression. Error bars represent ±1σ from three biological replicates. For each strain, an independent two-tailed t-test was used to compare the OD in the absence and presence of aTc (α=0.05). Significant differences are noted with asterisks.

To investigate if differences in Fd-dependent growth arise because each Fd is expressed at different levels, we created expression vectors that produce these proteins as fusions to red fluorescence protein (RFP). We performed whole cell fluorescence measurements using cells transformed with these vectors and compared the signal to cells expressing ml-Fd1 alone (Figure 7). This analysis revealed that pmN2-Fd is expressed at higher levels than the other three Fds under the expression conditions used for the cellular assay. This finding suggests that the weaker complementation observed with pmN2-Fd does not arise because it is expressed at lower levels than the Fds that complement growth.

**Figure 7.**
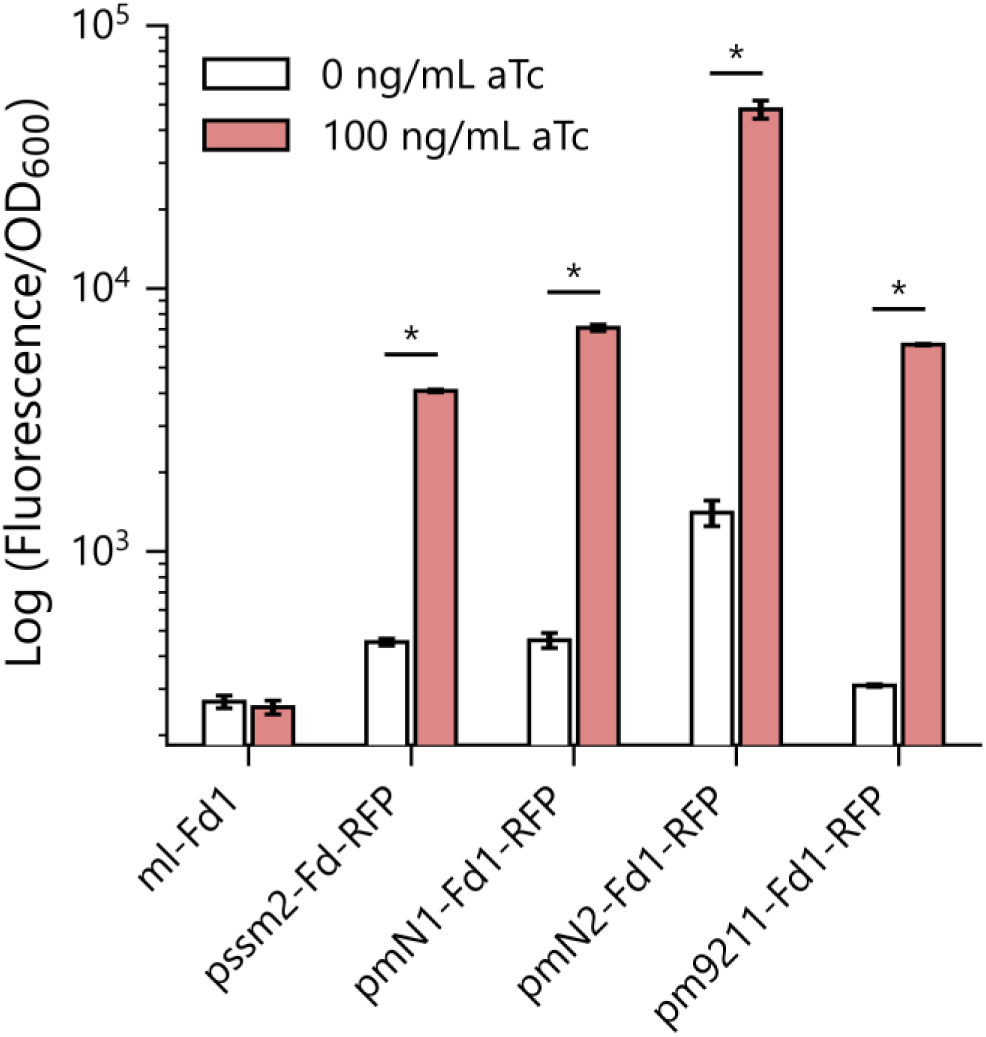
Expression of Fd-RFP fusion proteins. Fluorescence was measured in *E. coli* EW11 transformed with vectors that express Fds using an aTc-inducible promoter. Cells were grown in non-selective conditions. For each strain, an independent two-tailed t-test was used to compare fluorescence normalized to optical density (OD_600_) ±aTc (α=0.05). Error bars represent ±1σ from three biological replicates. Significant differences are noted with asterisks.

To evaluate whether variability in structural properties correlates with cellular complementation, we evaluated the surface charge distribution of Fd and SIR structures (zm-Fd1: 3B2F, pssm2-Fd: 6VJV, zm-SIR: 5H8V) as well as structural models for Fds and SIRs lacking structures (pmN2-Fd1, pm9211-Fd1, pmN2-SIR, and pm9211-SIR) (38, 71). With the Fds that complement corn SIR, we observed similar charge distribution proximal to the iron-sulfur cluster (Figure 8A). Notably, the host Fd that did not support cellular ET to corn SIR (pmN2-Fd1) is the least electronegative Fd. When we analyzed the charge distribution of the SIRs, we observed similar surface charges, with slightly more electronegative character in cyanobacterial SIRs compared to zm- SIR (Figure 8B).

**Figure 8.**
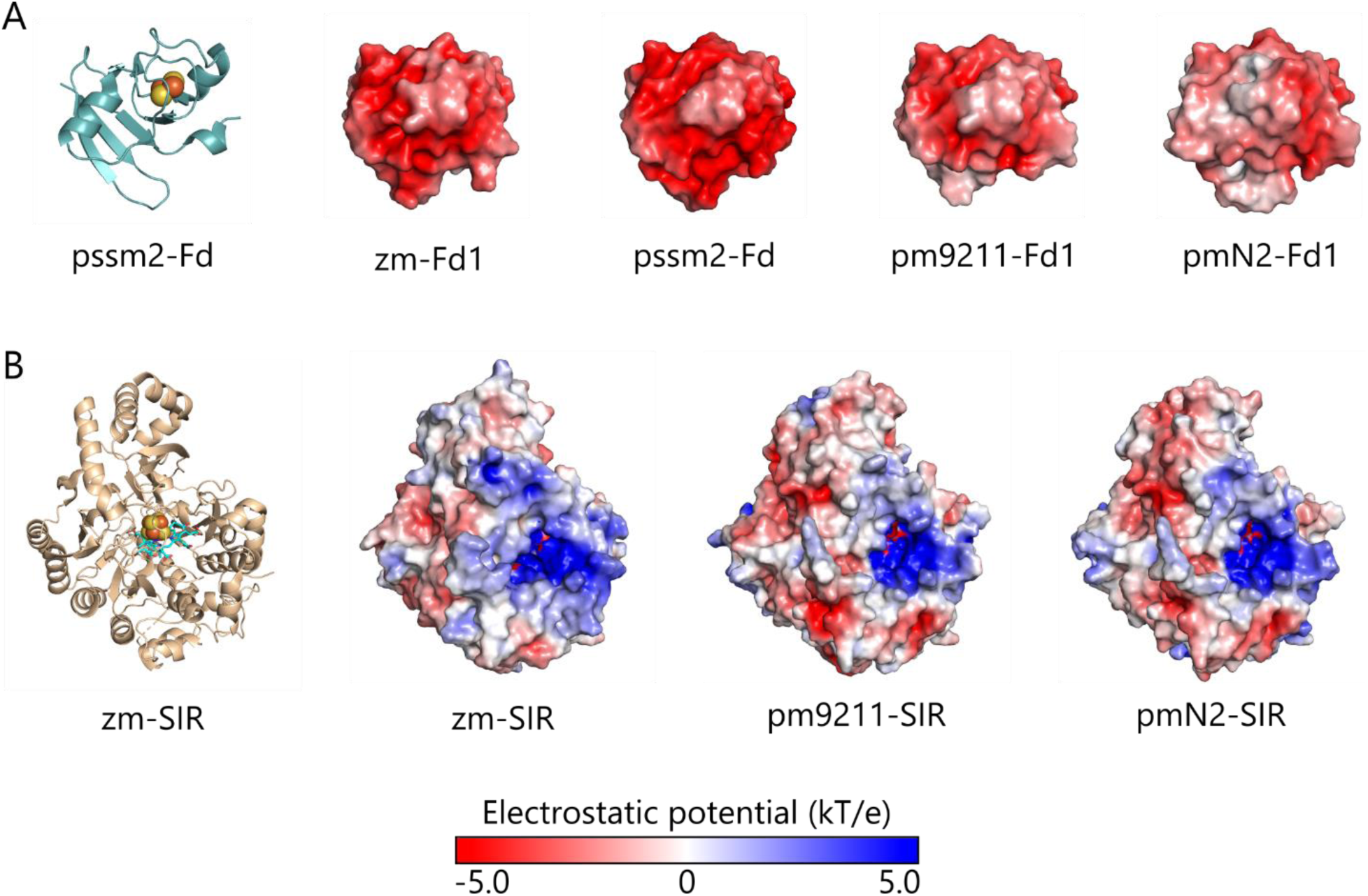
A comparison of the electrostatic potential surfaces of Fds and SIRs. The surfaces of (**A**) Fd and (**B**) SIR homologs are colored according to electrostatic potential, ranging from blue to red (kT/e), where kT is thermal energy, and e is the elementary charge. zm-Fd1, pssm2-Fd, and zm-SIR were generated using crystal structure data, while all other structures represent homology models (38, 70). The front faces of the Fds are negatively charged to varying degrees, while the SIRs display positive binding surfaces with small variations. These oppositely charged patches interact during binding, bringing the metallocofactors from each protein into close proximity to facilitate ET (38). Iron-sulfur clusters are displayed as spheres and sirohemes as sticks.

## Discussion

Our bioinformatic analysis reveals how phage Fd relates to the different Fd paralogs in cyanobacteria. Twelve major clusters of cyanobacterial Fds were observed, with some clusters representing Fds which are widespread across cyanobacteria, and others containing Fds from a small number of organisms. Only a small subset of the Fds in the network have been characterized. In the case of the cyanobacterial Fds in Cluster VI, which includes all of the cyanophage Fds, several of these proteins represent Fds expressed under phototrophic conditions (31, 74, 75). In some cases, this expression is likely due to their critical role as the electron acceptor for PSI (42, 45). Since Cluster VI contains cyanobacterial Fds that have evolved to support ET to a wide range of oxidoreductases, it seems likely that cyanophage Fds arose from the duplication of a cyanobacterial Fd that mediates ET to one or more of these partner oxidoreductases.

A comparison of biochemical studies in *Synechocystis* sp. PCC6803 with our bioinformatics suggests that the clusters represent Fd paralogs with distinct cellular roles. This organism has Fd paralogs that map onto Clusters V, VI, IX and X (31). The variable regulation of these Fds supports the idea that these proteins have distinct cellular roles (31, 74, 76). Transcripts encoding many of these Fd paralogs are regulated by environmental conditions, with changes in gene expression observed in response to salt stress, high metal concentrations, low light, and blue light. Despite this knowledge, we know little about the partner specificity profiles of Fds outside of Cluster VI.

Our bioinformatics revealed some additional trends across the clusters. Clusters I and II consist of mini- Fds with tight size distributions (µ = 79 ± 2, µ = 81 ± 5) and a tryptophan at the C-terminus. This is an unusual feature in comparison to canonical Fds in Cluster VI which generally end with a tyrosine. The tryptophan-terminating Fds are found in >40 cyanobacteria, although Cluster II consists wholly of *Prochlorococcus* and *Synechococcus* (10). Structures of these Fds have not yet been reported. The only structurally characterized [2Fe-2S] Fd with a terminal tryptophan is a cytochrome P450- reducing homolog from *Pseudomonas putida* (1PDX) but it is 106 amino acids long and much larger than the Fds within Cluster I and II Fds (77).

Some of the clusters in our network represent Fds that are unique to specific cyanobacterial species. Cluster III is solely populated by Fds from *P. marinus* ecotypes, which are not known to be infected by P-SSM2 cyanophage. Cluster V contains only Fds from *Synechocystis* strains. Finally, Cluster X is populated primarily by Fds from *P. marinus* and *Synechococcus sp*. strains, although it also contains two species of *Cyanobium*. Future studies will be required to better understand how sequence variation within and across clusters relates to the evolution of partner specificity profiles across different cyanobacteria.

Our biophysical analysis revealed that phage Fd exhibits a similar overall structure to non-host cyanobacterial Fds found in Cluster VI (41, 68–70), presenting ≤0.5 Å RMSD with mesophilic (nsp-Fd1 and s6803-Fd1) and thermophilic (te-Fd1 and ml- Fd1) Fds. Our electrochemical measurements showed that the pssm2-Fd has a potential on the high range of those observed in Cluster VI, which have midpoint potentials that range from -412 to - 336 mV (53, 78, 79). How the structures and electrochemical properties of *P. marinus* Fds relate to pssm2-Fd is not known. However, given the similarity of the phage Fd structure to more distantly-related Fds, it seems likely that they will be similar to characterized orthologs in other cyanobacteria that serve as PSI electron acceptors.

Our biochemical studies showed that pssm2-Fd has a low melting temperature (T_m_ = 29°C) compared with the other Fds in Cluster VI, which includes the thermophilic ml-Fd1 (80). This melting temperature is greater than the range (15 to 25°C) where the *P. marinus* hosts grow (4). This low T_m_ may have been selected to support more rapid turnover of cyanophage Fds or to regulate protein turnover in response to temperature changes. Currently, it is not known how Fds tune their stability without altering their midpoint potential. The native residues that control Fd stabilities (T_m_ = 29 to 76°C) could be established in future studies by creating chimeras between pssm2-Fd and more thermostable Fds with similar midpoint potentials and then evaluating how inheritance of different native residues relates to the thermostability of the resulting chimeras (81, 82).

Our cellular measurements suggest that cyanophages may need to support host sulfur metabolism to maximize their fitness. The underlying need for this ET is not known, but it may have arisen because sulfur-containing amino acids are required for translation of phage proteins under conditions where reduced sulfur is scarce. In *P. marinus* ecosystems, daytime dissolved dimethylsulfoniopropionate and methionine reach only 0.5-10 nM and 10-40 pM, respectively (83–85). In addition, methionine levels peak during the day and drop below detection limits after sunset suggesting that the diurnal cycle of cyanobacteria contributes to sulfur cycling in surface communities (84). In contrast, oxidized sulfur like sulfate is abundant, 28 mM (83), and it can be readily assimilated through a SIR-dependent pathway. Thus, cyanophages may use their Fd to support ET through this pathway under conditions where reduced sulfur becomes limiting to the production of phage proteins or other sulfur-containing metabolites critical to phage fitness.

The relative preference of phage Fd for host- versus phage-encoded oxidoreductases will be interesting to explore in future studies. One way to address this competition would be to examine how coexpression of the phage Fd-partner PebS and host SIR affects growth complementation in our cellular assay. The advantage of this synthetic biology approach over more traditional *in vitro* methods is that it would allow higher throughput analysis of protein-protein interactions in the crowded environment of a cell. Additionally, this approach could be applied to other natural and engineered Fds that support cellular ET from FNR to SIR (52, 85, 86).

## Experimental procedures

### Materials

Tris was from Fisher Scientific, N-cyclohexyl-3- aminopropanesulfonic (CAPS) was from Acros Organics, 2-(N-Morpholino)ethanesulfonic acid (MES) and N-[Tris(hydroxymethyl)methyl]-3- aminopropanesulfonic acid (TAPS) were from Fluka Biochemika, and isopropyl β-D-1- thiogalactopyranoside (IPTG), dithiothreitol (DTT), kanamycin, chloramphenicol, and streptomycin were from Research Product International. 3-Morpholinopropane-1-sulfonic acid (MOPS), N-Cyclohexyl-2- aminoethanesulfonic acid (CHES), and all other chemicals were purchased from Sigma-Aldrich. *E. coli* EW11 was a gift from Pam Silver (Harvard University) (72), *E. coli* XL1-Blue was from Agilent, and Rosetta(tm)(DE3) was from Novagen.

### Vector Design

All plasmids are listed in Table S3. Genes were synthesized by Integrated DNA technologies or Twist Bioscience as G-blocks and Gene Fragments, respectively. Plasmids were constructed by ligating PCR products amplified with Q5 polymerase (New England Biolabs) using Golden Gate assembly (87). Translation initiation sites were designed using the RBS calculator (88). All plasmids were sequence verified.

### Bioinformatics

To harvest viral Fd ORFs, all available viral genomes from Integrated Microbial Genomes and Microbiomes (n=8392) were scanned for genes encoding Fds with the Interpro signature IPR001041 (34). Viral genes with this IPR signature were compiled with the 807 cyanobacterial [2Fe-2S] Fds listed in Dataset S1. To obtain pairwise sequence similarity scoring and generate a SSN, these Fds were analyzed using the Enzyme Function Initiative-Enzyme Similarity Tool (36). The resulting networks were analyzed and images were generated using Cytoscape with the prefuse force directed layout based on an alignment score cutoff of 30 (89). Protein sequence alignments were generated using MUSCLE (90).

### Protein purification

*E. coli* Rosetta transformed with pJTA022 were grown at 37°C in lysogeny broth (LB) containing 50 µg/mL kanamycin to exponential phase, induced with 50 µM IPTG, and grown overnight at 37°C while shaking at 250 rpm. Cells harvested by centrifugation (4k x *g*) were resuspended in 25 mL lysis buffer (per L of culture), which contained 10 mM Tris pH 8, 5 mM DTT, 10 mg/L DNase I, and 0.5 mg/mL lysozyme. After freezing at −80°C, cells were thawed and mixed with a cOmplete Mini, EDTA-Free protease inhibitor tablet (Sigma-Aldrich) at a ratio of one tablet per 100 mL lysate. Samples were kept on ice or at 4°C for all subsequent purification steps. Clarified lysate generated by centrifugation (47k x *g*) was diluted three-fold with TED buffer (25 mM Tris pH 8, 1 mM EDTA, and 1 mM DTT) to lower the salt concentration. This mixture was loaded onto a DE52 column (Whatman), the column was washed with TED containing 200 mM NaCl, and the Fd was eluted using sequential isocratic washes with TED containing 250 and 300 mM NaCl. Fractions appearing brown were mixed and diluted with TED to bring NaCl below 100 mM. These pooled fractions were loaded onto HiTrap Q XL column (GE Healthcare) using an AKTA Start FPLC system (GE Healthcare). The column was washed using 100 mL TED buffer. Protein was eluted using a linear gradient (0 to 375 mM NaCl in TED) followed by an isocratic wash (500 mM NaCl in TED). Brown fractions were pooled and then further purified using a HiLoad 16/600 Superdex 75 column containing TED. SDS-PAGE was performed to analyze purity at each step using NuPage 12% Bis-Tris Gels (Invitrogen) and the PageRuler Unstained Broad Range Protein Ladder (Thermo Scientific). Samples appearing homogeneous were pooled and concentrated using an Amicon Ultra 10 K MWCO spin column and then flash frozen with liquid nitrogen. Following stability measurements, protein expression was evaluated at 25°C in Terrific Broth (TB). When protein was purified following low temperature expression, the final yield increased by 11.5-fold.

### Spectroscopy

Prior to all measurements, pssm2-Fd was dialyzed into TED. Absorbance spectra and ellipticity were acquired using a J-815 spectropolarimeter (Jasco, Inc.) using quartz cuvettes with a 1 cm path length. Scans used a 1 nm bandwidth, a 0.5 nm data pitch, and a 200 nm/min scan rate at 20°C. To assess stability, a cuvette with 50 µM pssm2-Fd was heated from 5 to 50°C at a rate of 1°C/min while monitoring ellipticity at 427 nm.

### Electrochemistry

Electrochemical measurements were performed anaerobically using a three-electrode system. A Ag/AgCl/1M KCl electrode (CH Instruments) was used as the reference electrode, and a platinum wire was used as the counter electrode. An edge-plane pyrolytic graphite electrode was used as the working electrode to perform protein film electrochemistry. Prior to adding Fd, this electrode was treated with 100 mM neomycin trisulfate (Sigma-Aldrich) to improve the electrochemical signal (91). An 3 µL aliquot of pssm2-Fd Fd (620 µM) was then applied to the electrode surface, and the protein was allowed to adhere for 1 min at 23°C prior to placing in a glass vial containing a pH 7 buffer solution (5 mM acetate, MES, MOPS, TAPS, CHES, CAPS) and 100 mM NaCl at 23.5 °C. Square wave voltammograms were collected at 10 Hz frequency, and electrochemical signals were analyzed using QSoas open software. Similar results were obtained when experiments were performed using pssm2-Fd from different purifications. A CH Instruments potentiostat and CHI660E electrochemical analyzer were used for all measurements. All data is reported relative to Standard Hydrogen Electrode (SHE), taking into account the potential difference between SHE and Ag/AgCl/1M KCl, which is 0.222 V.

### Crystallization

Purified pssm2-Fd (∼20 mg/mL) was used for crystal screening with a mosquitoLCP pipetting robot (SPT Labtech, Boston, MA) and commercially available screens including Wizard Classic 1 and 2 and Wizard Classic 3 and 4 (Rigaku Reagents, Inc., Bainbridge Island, WA), MORPHEUS and MIDAS (Molecular Dimensions, Holland, OH), and PEGRx and IndexHT (Hampton Research, Alisa Viejo, CA). Protein crystals were grown via the sitting drop method of vapor diffusion at a 1:1 (v:v) ratio of protein:precipitant 200:200 nL, at 20°C. Crystals for diffraction and data collection were optimized and grown with a precipitant solution containing 10% (w/v) PEG 3000, 200 mM zinc acetate, and 100 mM sodium acetate/acetic acid pH 4.5. The crystals were cryo- protected with 10% glycerol, harvested using a MiTeGen micromesh loop (MiTeGen, Ithaca, NY), and flash-cooled in liquid nitrogen prior to data collection.

### X-ray data collection, structure solution and refinement

Data were collected at the Advanced Photon Source GM/CA beamline 23ID-B using a 20 µm beam and 0.2 degree oscillations at a wavelength of 1.033 Å with an EIGER X 16M (DECTRIS Ltd, Philadelphia, PA) detector. The data were processed using the *autoPROC* toolbox (92), indexed and integrated using *XDS* (93), and scaled with *AIMLESS* (94). Initial phases were obtained by molecular replacement with *Phaser-MR* using the *Zea mays* Fd PDB ID: 5H57 (95) as a search model. Model building and refinement were performed with *Coot* (96) and *phenix*.*refine* (97). A custom graphics system was used for collaborative visualization (98). NCS restraints were used in initial refinements, with Chain A as a reference for Chain B, and removed in the final rounds of refinement. For TLS refinement, groups were automatically determined for Chain A while Chain B was treated as one group. Anomalous difference maps were calculated using *ANODE* and zinc from the precipitant solution was modeled in the structure (99). Data processing and refinement software were compiled and supported by the SBGrid Consortium (100). The structural model is available in the worldwide Protein Data Bank (PDB ID: 6VJV) (101). Data collection and refinement statistics are provided in Table S2.

### Homology modeling

All-atom homology models were generated for pmN2-Fd1, pm9211-Fd1, pmN2-SIR, and pm9211-SIR using the fold and function assignment system (FFAS) server (102, 103). The Fe-S clusters for the Fd homology models were incorporated by aligning and copying them from our experimental model, PDB: 6VJV. The Fe-S clusters and sirohemes for the SIR homology models were incorporated by aligning the models to PDB: 5H8V and copying them.

### Structural analysis

Root-mean-squared deviations were calculated, hydrogen bonds were detected (length cutoff: 3.5 Å), and structural images were generated using PyMOL (104). The illustration of conserved Fd residues mapped onto the pssm2-Fd structure was generated using the ConSurf Server with the WAG evolution model, seeded with all Fd sequences from Cluster VI (71). Electrostatic surface representations were calculated in PyMOL (version 2.3.2) using the APBS electrostatics plug-in with the prepwizard method default (Schrödinger) (104). All solvent molecules, ions, and ligands were removed, while the Fe-S clusters and sirohemes were used in the map calculation.

### Cellular assay

To assess Fd ET from FNR to SIR, *E. coli* EW11 was transformed with two plasmids using electroporation as described (52, 72, 85, 86). One plasmid constitutively expresses FNR and SIR pairs, while the other expresses Fds under control of an anhydrotetracycline (aTc)-inducible promoter. To maintain both plasmids, all growth steps included chloramphenicol (34 μg/mL) and streptomycin (100 μg/mL). Starter cultures inoculated using single colonies were grown in deep-well 96-well plates for 18 h at 37°C in 1 mL of a nonselective modified M9 medium (M9c), which contained sodium phosphate heptahydrate, dibasic (6.8 g/L), potassium phosphate, monobasic (3 g/L), sodium chloride (0.5 g/L), 2% glucose, ammonium chloride (1 g/L), calcium chloride (0.1 mM), magnesium sulfate (2 mM), ferric citrate (500 μM), p-aminobenzoic acid (2 mg/L), inositol (20 mg/L), adenine (5 mg/L), uracil (20 mg/L), tryptophan (40 mg/L), tyrosine (1.2 mg/L), and the remaining 18 amino acids (80 mg/L each). Starter cultures grown to stationary phase were then diluted 1:100 into a selective modified M9 medium (M9sa), which is identical to M9c but lacks cysteine and methionine. Cultures were inoculated in sterile Nunc(tm) Edge 2.0 96-well plates with side reservoirs filled with 1 mL of water. The edge between plate tops and bottoms were sealed with parafilm. Cells were grown in a Spark plate reader (Tecan) at 37°C while shaking (90 rpm) at an amplitude of 3 mm in double-orbital mode. Optical density (OD) was measured at 600 nm every 5 min for 48 h. For endpoint measurements, all conditions were the same except cells were grown in incubators at the indicated temperature while shaking at 250 rpm for 48 hours.

### Fluorescence measurements

To evaluate the cellular expression of Fds, the genes encoding each Fd were expressed as fusions to a monomeric RFP with a twelve amino acid linker (105). Each vector was transformed into *E. coli* EW11 cells, alongside a plasmid (pSAC01) constitutively expressing *Zea mays* SIR (zm-SIR) and FNR (zm-FNR) (52). Transformed cells were grown under identical conditions to those described for the cellular assay, with the exception of being grown in M9c at all steps. After 24 hours, endpoint OD and RFP fluorescence (λ_excitation_ = 560 nm; λ_emission_ = 650 nm) were measured.

### Statistics

Error bars represent standard deviation calculated from three or more biological replicates. Independent two-tailed t-tests were applied to compare differences between all relevant samples with α=0.05.

## Data availability

The structure presented in this study is available in the PDB with the code 6VJV. All remaining data are contained within this article.

## Funding and additional information

This project was supported by DOE grant DE-SC0014462 (to J.J.S), NASA NAI grant 80NSSC18M0093 (to G.N.B and J.J.S), NSF grant 1231306 (to G.N.P), and Moore Foundation grant 7524 (to G.N.B and J.J.S.). GM/CA@APS has been funded in whole or in part with Federal funds from the National Cancer Institute (ACB-12002) and the National Institute of General Medical Sciences (AGM-12006). The Eiger 16M detector at GM/CA-XSD was funded by NIH grant S10 OD012289. This research used resources of the Advanced Photon Source, a DOE Office of Science User Facility operated for the DOE Office of Science by Argonne National Laboratory under Contract No. DE- AC02-06CH11357. I.J.C. and J.T.A. were supported by a Lodieska Stockbridge Vaughn Fellowship.

## Author contributions

I.J.C., J.T.A., G.N.B., and J.J.S. conceptualized the project. I.J.C. and J.L.O. performed bioinformatics. I.J.C., N.S., and J.T.A. purified recombinant protein. I.J.C. conducted spectroscopy. J.L.O., W.X., M.D.M., and G.N.P. crystallized pssm2-Fd and generated the structures. D.K. conducted electrochemistry. I.J.C. constructed vectors and performed cellular assays. I.J.C. and J.J.S. wrote the manuscript, and all other authors refined the text.

## ABBREVIATONS

aTc: anhydrotetracycline
DTT: dithiothreitol
ET: electron transfer
Fd: ferredoxin
FNR: Fd-NADP reductase
GcpE: 4-Hydroxy-3-methylbut-2-enyl diphosphate synthase
GS: glutamate synthase
HO: heme oxygenase
IPTG: isopropyl β-D-1-thiogalactopyranoside
LB: lysogeny broth
ml-Fd1: *Mastigocladus laminosus* Fd 1
NIR: nitrite reductase
nsp-Fd1: *Nostoc sp*. PCC 7119 Fd 1
OD: optical density
ORFs: open reading frames
PcyA: phycocyanobilin:Fd oxidoreductase
PebS: phycoerythrobilin synthase
pm9211-Fd1: *Prochlorococcus marinus* MIT9211 Fd 1
pm9211-SIR: *Prochlorococcus marinus* MIT9211 SIR
pmN1-Fd1: *Prochlorococcus marinus* NATL1A Fd 1
pmN2-Fd1: *Prochlorococcus marinus* NATL2A Fd 1
pmN2-SIR: *Prochlorococcus marinus* NATL2A SIR
PSI: photosystem I
pssm2-Fd: myovirus P-SSM2 Fd
RFP: red fluorescent protein
s6803-Fd1: *Synechocystis sp*. PCC 6803 Fd 1
SHE: standard hydrogen electrode
SIR: sulfite reductase
SSN: sequence similarity network
te-Fd1: *Thermosynechococcus elongatus* Fd 1
zm-Fd1: *Zea mays* Fd 1
zm-FNR: *Zea mays* FNR
zm-SIR: *Zea mays* SIR

