## Supporting Information for "*Prochlorococcus* phage ferredoxin: structural characterization and electron transfer to cyanobacterial sulfite reductases"

**Running title:** *Phage Fd characterization and host SIR interactions*

---

**Supporting Information**

***List of materials***

1. Supplementary Table 1
2. Supplementary Table 2
3. Supplementary Table 3
4. Supplementary Figure 1
5. Supplementary Figure 2
6. Supplementary Figure 3
7. Supplementary Dataset 1 (separate file)

**Table S1. Data collection and refinement statistics.**

| PDB ID: 6VJV | |
|---|---|
| # of copies in asymmetric unit | 2 |
| Space group | $P2_1$ |
| Unit cell parameters (Å) | a= 52.62 b= 31.45 c= 61.29, β= 92.49 |
| Wavelength (Å) | 1.033 |
| Data collection statistics | |
| Resolution range (Å) | 52.6 – 1.59 (1.59 – 1.62)[a] |
| Number of reflections | 178386 (5968) |
| Completeness (%) | 98.5 (86.1) |
| $R_{merge}$[b] | 0.11 (1.77) |
| Redundancy | 6.60 (5.07) |
| Mean I/σ(I) | 8.9 (1.0) |
| Resolution <I/σ(I)> = 2 (Å) | 1.68 |
| $CC_{1/2}$[c] | 0.996 (0.319) |
| Refinement statistics | |
| Resolution range (Å) | 39.06 – 1.59 |
| $R_{cryst}$[d]/$R_{free}$[e] | 0.180/0.213 |
| RMSD bonds (Å) | 0.02 |
| RMSD angles (deg) | 1.56 |
| Average B factor ($Å^2$) | 32.5 |
| Number of water molecules | 212 |
| Number of acetate molecules | 4 |
| Number of zinc molecules | 14 |
| Ramachandran favored (%) | 97.34 |
| Ramachandran allowed (%) | 2.66 |
| FES RSCC (Chain A / Chain B) | 0.99 / 0.94 |

[a]Values in parentheses are for the highest-resolution shell.
[b]$R_{merge} = \Sigma_h \Sigma | I_i(h) - \langle I(h) \rangle | / \Sigma_h \Sigma_i I_i(h)$, where $I_i(h)$ is the intensity of an individual measurement of the reflection and <I(h)> is the mean intensity of the reflection.
[c]$CC_{1/2} = \sum (x - \langle x \rangle)(y - \langle x \rangle)/[\sum(x - \langle x \rangle)^2 \Sigma(y - \langle y \rangle)^2]^{1/2}$
[d]$R_{cryst} = \Sigma_h | |F_{obs}| - |F_{calc}| | / \Sigma_h |F_{obs}|$, where $F_{obs}$ and $F_{calc}$ are the observed and calculated structure factor amplitudes, respectively.
[e]$R_{free}$ was calculated as $R_{cryst}$ using 5% of the randomly selected unique reflections that were omitted from structure refinement.

**Table S2. Structural comparison of pssm2-Fd and cyanobacteria homologs.** RMSD of cyanophage pssm2-Fd and other cyanobacterial Fds quantified in PyMOL. Salt bridges are defined as an interaction between positively and negatively charged amino acids within 4.0 Å. Hydrogen bonds are defined as interactions between hydrogen-bond-donating atoms and hydrogen-bond-accepting atoms within 3.5 Å, and these are further subdivided into intramolecular hydrogen bonds versus those with the solvent.

| Species | Fd name | PDB ID | RMSD with pssm2-Fd (Å) | Salt Bridges | Hydrogen Bonds (intramolecular + w/ solvent) |
|---|---|---|---|---|---|
| *Prochlorococcus* phage P-SSM2 | pssm2-Fd | 6VJV | - | 3 | 133 (76 + 57) |
| *Synechocystis sp.* PCC 6803 | s6803-Fd1 | 1OFF | 0.433 | 5 | 115 (67 + 48) |
| *Thermosynechococcus elongatus* | te-Fd1 | 5AUI | 0.501 | 4 | 105 (80 + 25) |
| *Nostoc sp*. PCC 7119 | nsp-Fd1 | 1CZP | 0.472 | 4 | 154 (80 + 74) |
| *Mastigocladus laminosus* | ml-Fd1 | 1RFK | 0.391 | 5 | 123 (67 + 56) |
| *Zea mays* | zm-Fd1 | 3B2F | 0.455 | 6 | 107 (73 + 34) |

**Table S3. Vectors constructed.** Vectors constructed in this study are listed with antibiotic markers, plasmid origin, and genes expressed.

| Plasmid Name | Addgene ID | Description |
|---|---|---|
| pSAC01 | 131826 | $Spec^R$, p15a vector constitutively expressing *Zea mays* FNR and *Z. mays* SIR |
| pSAC16 | 137965 | $Spec^R$, p15a vector constitutively expressing *Zea mays* FNR and *Prochlorococcus marinus* NATL2A SIR |
| pSAC17 | 137966 | $Spec^R$, p15a vector constitutively expressing *Zea mays* FNR and *Prochlorococcus marinus* MIT9211 SIR |
| pFd007 | 131828 | $Cam^R$, ColE1 vector with aTc inducible Mastigocladus laminosus Fd1 |
| pFd022.R2 | 137967 | $Cam^R$, ColE1 vector with aTc inducible *Prochlorococcus* phage P-SSM2 Fd |
| pFd029.R2 | 137968 | $Cam^R$, ColE1 vector with aTc inducible *Prochlorococcus marinus* NATL1A Fd1 |
| pFd030.R2 | 137969 | $Cam^R$, ColE1 vector with aTc inducible *Prochlorococcus marinus* NATL2A Fd1 |
| pFd031 | 137970 | $Cam^R$, ColE1 vector with aTc inducible *Prochlorococcus marinus* MIT9211 Fd1 |
| pFd022.R2.RFP | 137971 | $Cam^R$, ColE1 vector with aTc inducible *Prochlorococcus* phage P-SSM2 Fd-L12-RFP |
| pFd029.R2.RFP | 137972 | $Cam^R$, ColE1 vector with aTc inducible *Prochlorococcus marinus* NATL1A Fd1-L12-RFP |
| pFd030.R2.RFP | 137973 | $Cam^R$, ColE1 vector with aTc inducible *Prochlorococcus marinus* NATL2A Fd1-L12-RFP |
| pFd031.RFP | 137974 | $Cam^R$, ColE1 vector with aTc inducible *Prochlorococcus marinus* MIT9211 Fd1-L12-RFP |
| pJTA022 | 137975 | $Kan^R$, ColE1 pET28b derived vector with T7-lac inducible *Prochlorococcus* phage P-SSM2 Fd |

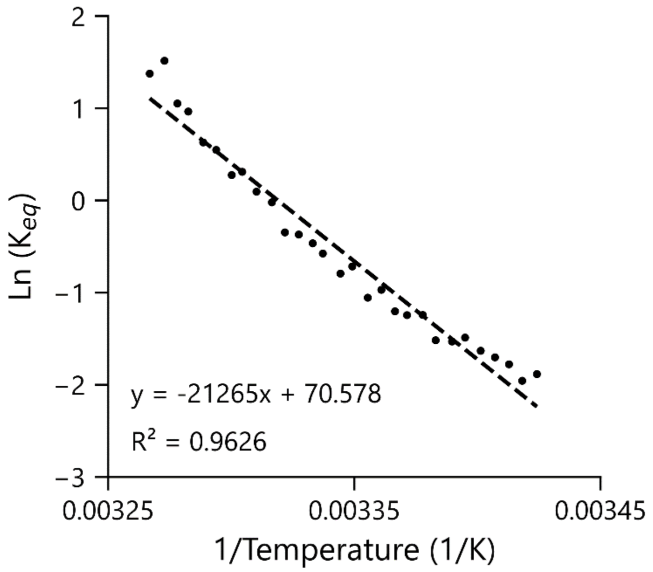


**Figure S1. Van't Hoff analysis of pssm2-Fd melting curve.** A plot of ln $K_{eq}$ vs. 1/T for the melting of pssm2-Fd (50 µM) measured by 427 nm ellipticity. With a linear fit closest to the transition (0.00325 to 0.00345 1/K), the slope yields $\Delta H_{unfolding}$ as 177 kJ·mol$^{-1}$ and the y-intercept yields $\Delta S_{unfolding}$ as 587 J·mol$^{-1}$·K$^{-1}$, when both are multiplied by the ideal gas constant *R* (8.314 J·mol$^{-1}$·K$^{-1}$). Assuming ΔG=0 at the folding transition, the $T_m$ is calculated by the ratio of $\Delta H_{unfolding}/\Delta S_{unfolding}$, yielding $T_m$ =301 K (28°C).

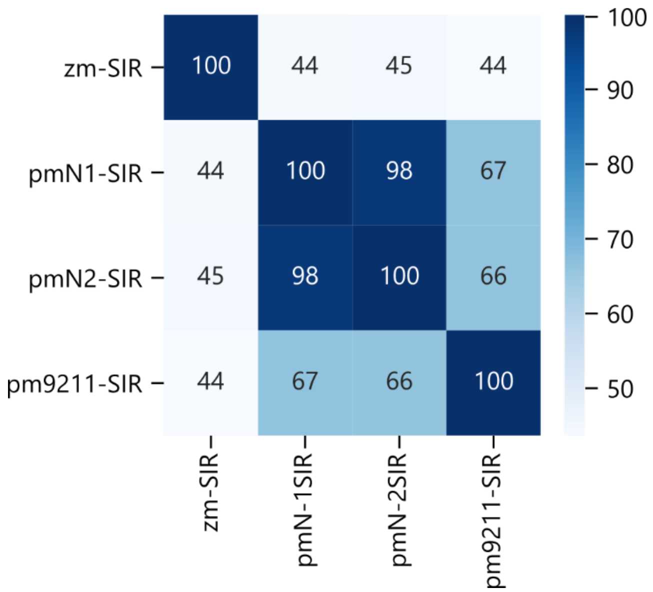


**Figure S2. Pairwise sequence identities of *Prochlorococcus marinus* homologs.** Sequence identity matrix of *Prochlorococcus marinus* SIRs and zm-SIR, with percentages rounded to the nearest integer. Positions corresponding to more closely related SIRs are shown in darker shades of blue. Abbreviations used: zm-SIR (*Zea mays* sulfite reductase), pmN1-SIR (*P. marinus* NATL1A sulfite reductase), pmN2-SIR (*P. marinus* NATL2A sulfite reductase), and pm9211-SIR (*P. marinus* MIT9211 sulfite reductase).

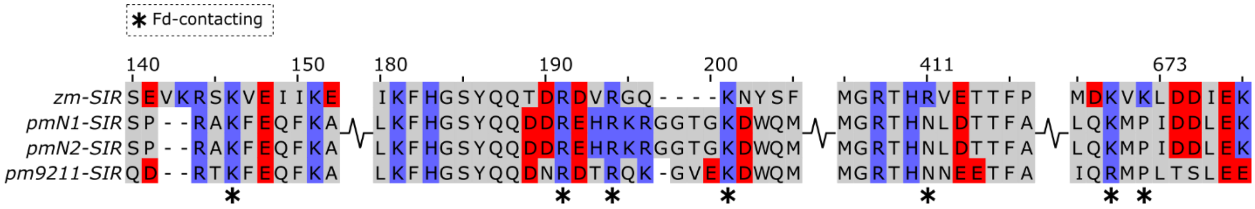


**Figure S3. Host cyanobacterial SIR sequence comparisons.** A multiple sequence alignment illustrates the conservation of charged residues and residues implicated in binding Fds (asterisks). Negatively charged residues are colored red and positively charged residues are colored blue. Abbreviations used: zm-SIR (*Zea mays* sulfite reductase), pmN1-SIR (*P. marinus* NATL1A sulfite reductase), pmN2-SIR (*P. marinus* NATL2A sulfite reductase), and pm9211-SIR (*P. marinus* MIT9211 sulfite reductase).

**Dataset S1. List of cyanobacterial and cyanophage sequences used to generate Figure 1 (*provided in separate file*).** The sequences harvested from the JGI/IMG database are ordered according to cluster location, alongside information about their clusters, including Gene ID, Species, Clade, Sequence Length, Cluster Number, Cluster Species Count, Cluster Average Length, Cluster Length Standard Deviation, and Sequence.
